# The IncC and IncX1 resistance plasmids present in multi-drug resistant *Escherichia coli* strains isolated from poultry manure in Poland

**DOI:** 10.1101/2024.04.05.588246

**Authors:** Magdalena Zalewska, Aleksandra Błażejewska, Jan Gawor, Dorota Adamska, Krzysztof Goryca, Michał Szeląg, Patryk Kalinowski, Magdalena Popowska

## Abstract

The study describes the whole-genome sequencing of two antibiotic-resistant representative *Escherichia coli* strains, isolated from poultry manure in 2020. The samples were obtained from a commercial chicken meat production facility in Poland. The antibiotic resistance profile was characterized by co-resistance to β-lactam antibiotics, aminoglycosides, and fluoroquinolones. The three identified resistance plasmids (R-plasmids), pECmdr13.2, pECmdr13.3 and pECmdr14.1, harbored various genes conferring resistance to tetracyclines (*tetR[A]*) for, aminoglycoside (*aph, aac* and *aad* families), β-lactam (*bla_CMY-2_*, *bla_TEM-176_*), sulfonamide (*sul1, sul2*), fluoroquinolone (*qnrS1*), and phenicol (*floR*). These plasmids, which have not been previously reported in Poland, were found to carry IS26 insertion elements, the intI1-integrase gene, as well as conjugal transfer genes, facilitating horizontal gene transfer. Plasmids pECmdr13.2 and pECmdr14.1 also possessed a mercury resistance gene operon related to transposon Tn1696; this promotes plasmid persistence even without antibiotic selection pressure due to co-selection mechanisms such as co-resistance. The chicken manure-derived plasmids belonged to the IncX1 (narrow host range) and IncC (broad host range) incompatibility groups. Similar plasmids have been identified in various environments, clinical isolates, and farm animals, including cattle, swine, and poultry. This study holds significant importance for the One Health approach, as it highlights the potential for antimicrobial-resistant bacteria from livestock and food sources, particularly *E. coli*, to transfer through the food chain to humans and vice versa. This underscores the need for vigilant monitoring of R-plasmids prevalence in the human, animal and natural environments, and to implement strategies to mitigate the spread of antibiotic resistance.

## Introduction

Years of misuse and overuse of antibiotics in the livestock and poultry industry have resulted in many irreversible changes, not only in the animal production sector but also in the everyday life of each human. Extensive, uncontrolled antibiotic use results in the presence of low, sub-inhibitory concentrations in the tissues and guts of treated animals (1) and the environment (2). The mechanisms of action of sublethal levels of antibiotics on bacteria are described in detail by Andersson and Hughes (3). In food-producing animals, antibiotic use affects the functions of enteric bacteria and can temporarily increase antibiotic resistance in the fecal microbiome (4, 5). The chronic application of such sub-therapeutic doses favors the selection of antibiotic-resistant bacteria (ARB) by promoting their growth or introducing *de novo* mutations. The resistance mechanism of a particular ARB depends on its harbored antibiotic-resistance genes (ARGs). The majority of ARG transmission occurs *via* horizontal gene transfer (HGT), whereby mobile genetic elements (MGEs), such as plasmids or transposons coding for ARG, are exchanged between bacterial species, even those not closely related (6).

*Escherichia coli* is classified as a rod-shaped, non-sporulating, facultatively anaerobic Gram-negative bacteria belonging to the family *Enterobacteriaceae*. This species mainly lives in the lower intestinal tract of warm-blooded animals, including livestock and poultry, and is often excreted into the environment with urine and feces. In addition to being a widespread gut commensal of vertebrates, *E. coli* is also a versatile pathogen, killing more than two million humans annually through intraintestinal and extraintestinal diseases (7). The presence of *E. coli* in the natural environment has long been considered an indicator of recent fecal contamination (8), but many studies show that many strains can survive and proliferate in such conditions and be incorporated into indigenous microbial communities (9, 10).

The presence of *E. coli* and other commensal bacteria in the animals’ gut can be beneficial. Some help animals to digest complex molecules, which cannot otherwise be assimilated, thus producing vitamin K or certain B vitamins essential for the host; others prevent gut colonization by competing with harmful microorganisms, i.e. by consuming available nutrients, occupying physical space, and producing substances such as bacteriocins that inhibit the growth of the pathogen. They can also play an essential role in the development and regulation of the immune system by stimulating it and helping it to differentiate between harmful and non-harmful organisms (7, 11, 12).

However, some strains of *E. coli* can be pathogenic, causing severe diseases in humans and animals. Infection with pathogenic *E. coli* strain can manifest in enteric or diarrheal disease, urinary tract infections, and meningitis or sepsis. The intestinal symptoms may be caused by EIEC (enteroinvasive *E. coli),* EHEC (enterohaemorrhagic *E. coli*), ETEC (enterotoxigenic *E. coli*), EPEC (enteropathogenic *E. coli*), DAEC (diffusely adherent *E. coli*), and EAEC (enteroaggregative *E. coli*) (13). The most common extraintestinal *E. coli* infection is urinary tract infection, caused by UPEC (uropathogenic *E. coli*); however the pathotype MNEC (meningitis-associated *E. coli*), responsible for meningitis and sepsis, is becoming increasingly prevalent. EPEC, EHEC, and ETEC can also cause animal disease using the same virulence factors present in human strains and colonization factors unique to animals. The avian pathogenic *E. coli* (APEC) is an additional animal pathotype, causing extraintestinal infections of poultry, primarily respiratory infections, pericarditis, and septicemia (13).

The aim of the study was to examine the population of *E. coli* strains isolated from poultry (from intensive rearing facilities) feces. Based on the results of antimicrobial susceptibility testing and plasmid profile characteristics, the most prevalent strains were subjected to sequencing with NovaSeq 6000 (Illumina) and MinIon (Oxford Nanopore) platforms for full plasmid structure recovery. Two representative multi-drug resistance (MDR) *E. coli* isolates, were found to harbour three resistance plasmids (R-plasmids) that had not previously been reported in Poland: pECmdr13.3 (IncX1), pECmdr13.2 and pECmdr14.1 (IncC). These were found to carry genes coding for resistance to tetracyclines, aminoglycosides, β-lactams, sulfonamides, fluoroquinolones, and phenicol; the last two were accompanied by mercury resistance.

## Results

### *E. coli* strain selection

Selective media were used to isolate 62 bacterial strains exhibiting growth characteristic of *E. coli*: 51 strains were identified on Eosin Methylene Blue (EMB) Agar and 11 on MacConkey Agar. Of all isolated bacteria, 23 were identified as *E. coli* by MALDI-TOF MS/MS analysis: of these, 20 strains were isolated on EMB agar, three on MacConkey agar, 13 from chicken manure (CM), and 10 from chicken litter (CL). All bacteria strains were isolated on agar plates supplemented with cefotaxime. No strains resistant to imipenem were identified. All strains were subjected to antimicrobial susceptibility profile determination by Vitek 2 Compact: the results are listed in S1 Table 2. Isolates exhibiting similar antimicrobial susceptibility profiles were grouped, and their plasmidic profiles were determined. After comparing the plasmidic profiles within the groups, four potentially unique bacterial strains were chosen for plasmid structure determination by total DNA sequencing.

### Genomes sequencing and assembly

High-quality genome sequence assemblies for the selected strains were obtained by combining scaffolds from Oxford Nanopore long-read technology (MINIon platform) with Illumina short-read technology (2×100 nt in pair-end mode, NovaSeq 6000, approximate 100x coverage) for quality improvement. The chromosome was successfully assembled, resulting in close to 5Mb contigs for each strain, and three additional contigs over 40 kb were reconstructed.

### Plasmids

Six plasmids belonging to two bacterial isolates were identified (Table 1.).

**Table 1.** Plasmids identified in studied *E. coli* strains. Plasmids harboring ARGs are marked in gray.

| strain no. | origin | plasmid no. | plasmid name in our collection | size [bp] | Accession no. |
| --- | --- | --- | --- | --- | --- |
| 13 | chicken manure | 1 | pECmdr13.1 | 174994 |  |
|  |  | 2 | pECmdr13.2 | 158923 |  |
|  |  | 3 | pECmdr13.3 | 45794 |  |
| 14 | chicken manure | 1 | pECmdr14.1 | 297035 |  |
|  |  | 2 | pECmdr14.2 | 157029 |  |
|  |  | 3 | pECmdr14.3 | 99355 |  |

Three of the identified plasmids harbored different ARGs and MGEs (Tab. 2). The linear structure of the plasmid structures, highlighting the ARGs, MGEs, conjugal transfer genes, transposon transposase genes, heavy metal resistance genes, replication system, integron integrase, partitioning system elements, is presented in Figure 1-3. A detailed analysis inidicated close similarity between the identified plasmids and some bacterial plasmid sequences deposited in the PLSDB plasmid database (https://ccb-microbe.cs.uni-saarland.de/plsdb/); however, there were some differences between them. Plasmid pECmdr13.2 has 133 similar hits, isolated from varied sources located in different regions, plasmid pECmdr13.3 has 18 similar hits, and pECmdr14.1 has 155 similar hits (identity threshold 0.99). None of these plasmids has previously been reported in Poland, but they have been identified in many countries. Although plasmids similar to pECmdr13.2, pECmdr14.1 have been found in the environment, clinical isolates, and farm animals (cattle, swine, and poultry/birds), plasmids similar to pECmdr13.3 have been identified in the same places but only from poultry. Plasmids that do not confer antibiotic resistance will be characterized and deposited further as a part of an additional project.

**Figure 1.**
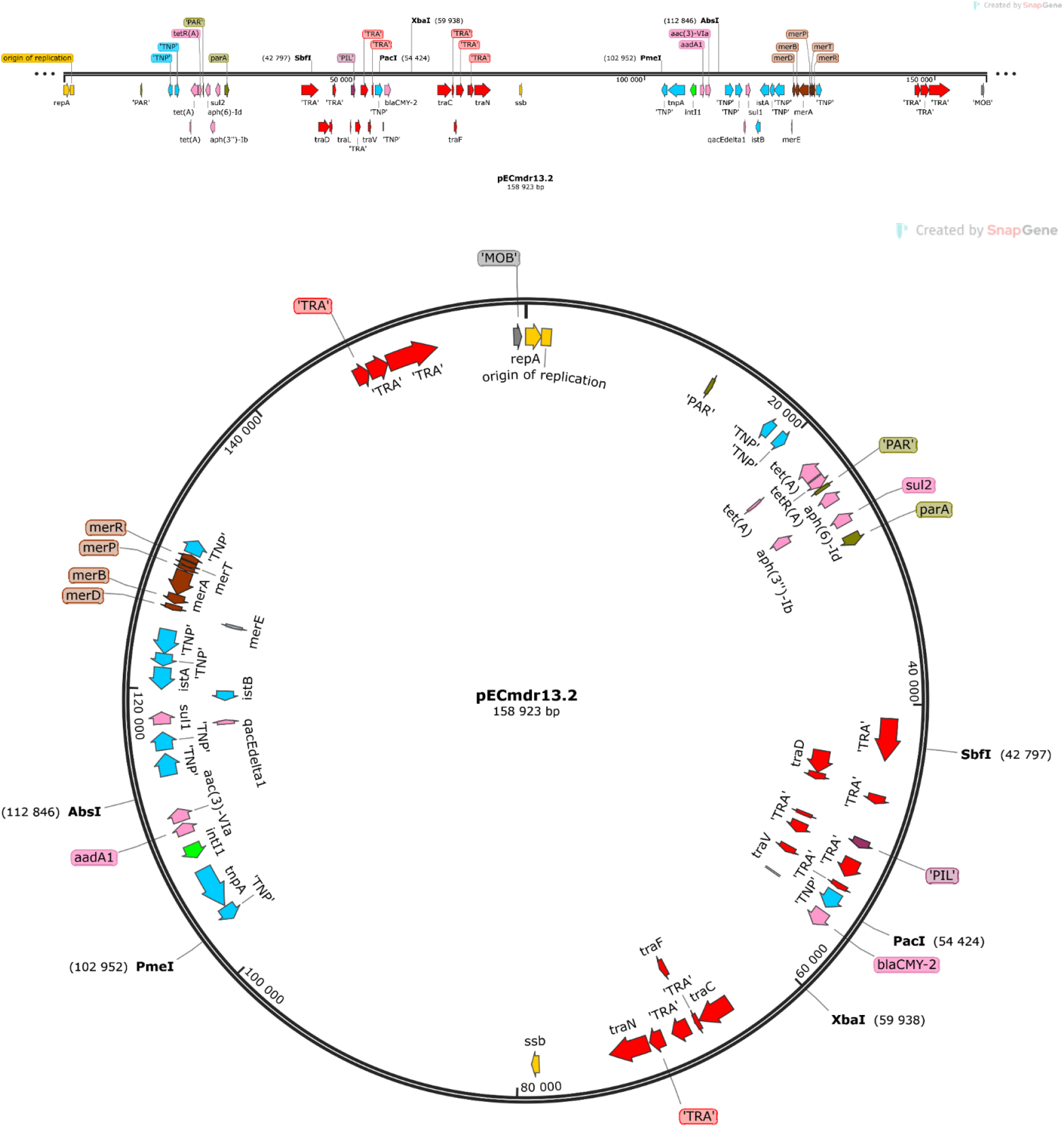
Linear and circular representation of pECmdr13.2 plasmid; pink-ARGs, red-conjugal transfer genes (TRA), blue-transposon transposase genes (TNP), brown-heavy metal resistance genes, yellow-replication system (REP), green-integron integrase, khaki-partitioning system PAR, gray-mob system (MOB), plum-pilus system (PIL); a name written in quotation marks indicates that the gene function has been assigned based on the homology of the protein it encodes

**Figure 2.**
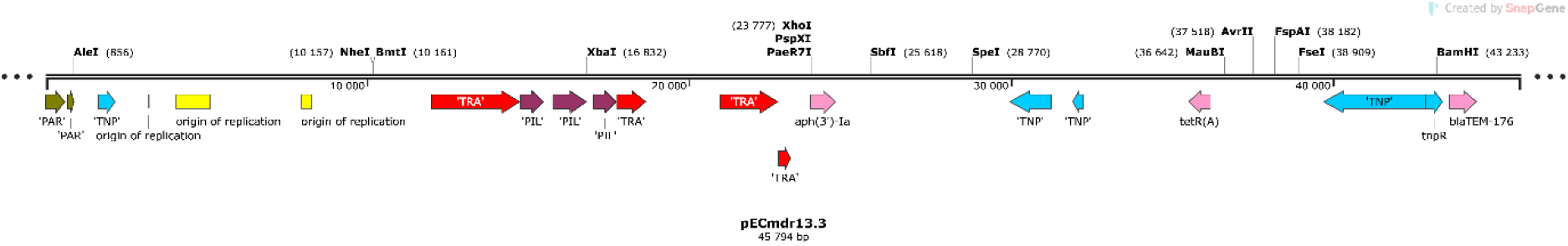
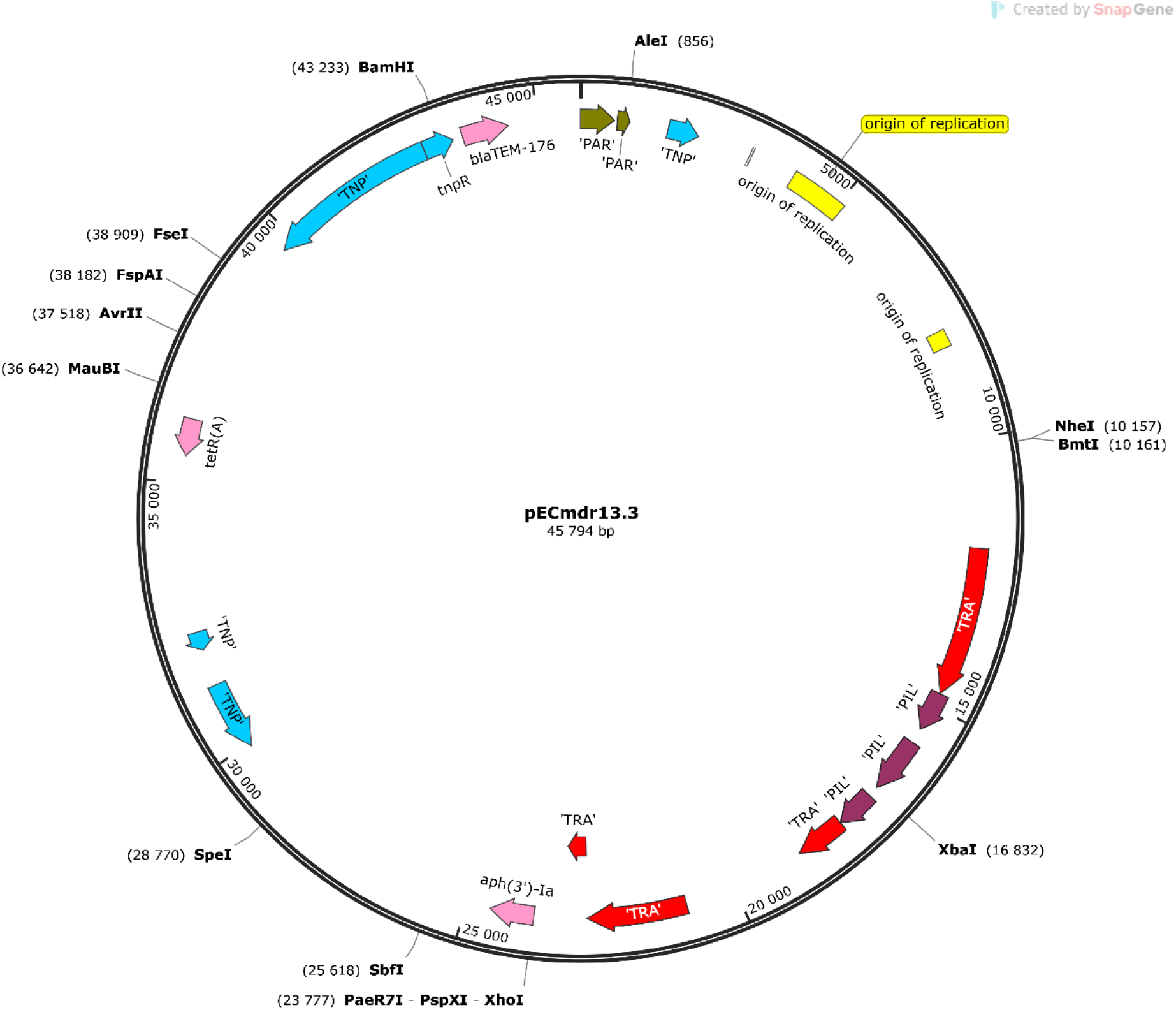
Linear and circular representation of pECmdr13.3 plasmid; pink-ARGs, red-conjugal transfer genes (TRA), blue-transposon transposase genes (TNP), brown-heavy metal resistance genes, yellow-replication system (REP), green-integron integrase, khaki-partitioning system (PAR), gray-mob system (MOB), plum-pilus system (PIL); a name given in quotation marks indicates that the gene function has been assigned based on the homology of the protein it encodes

**Figure 3.**
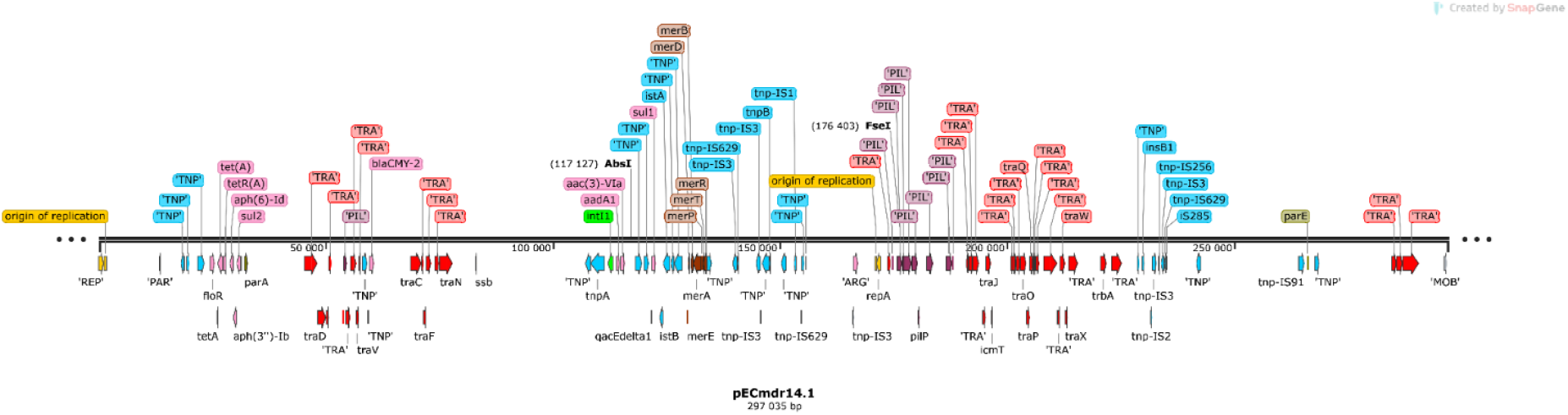
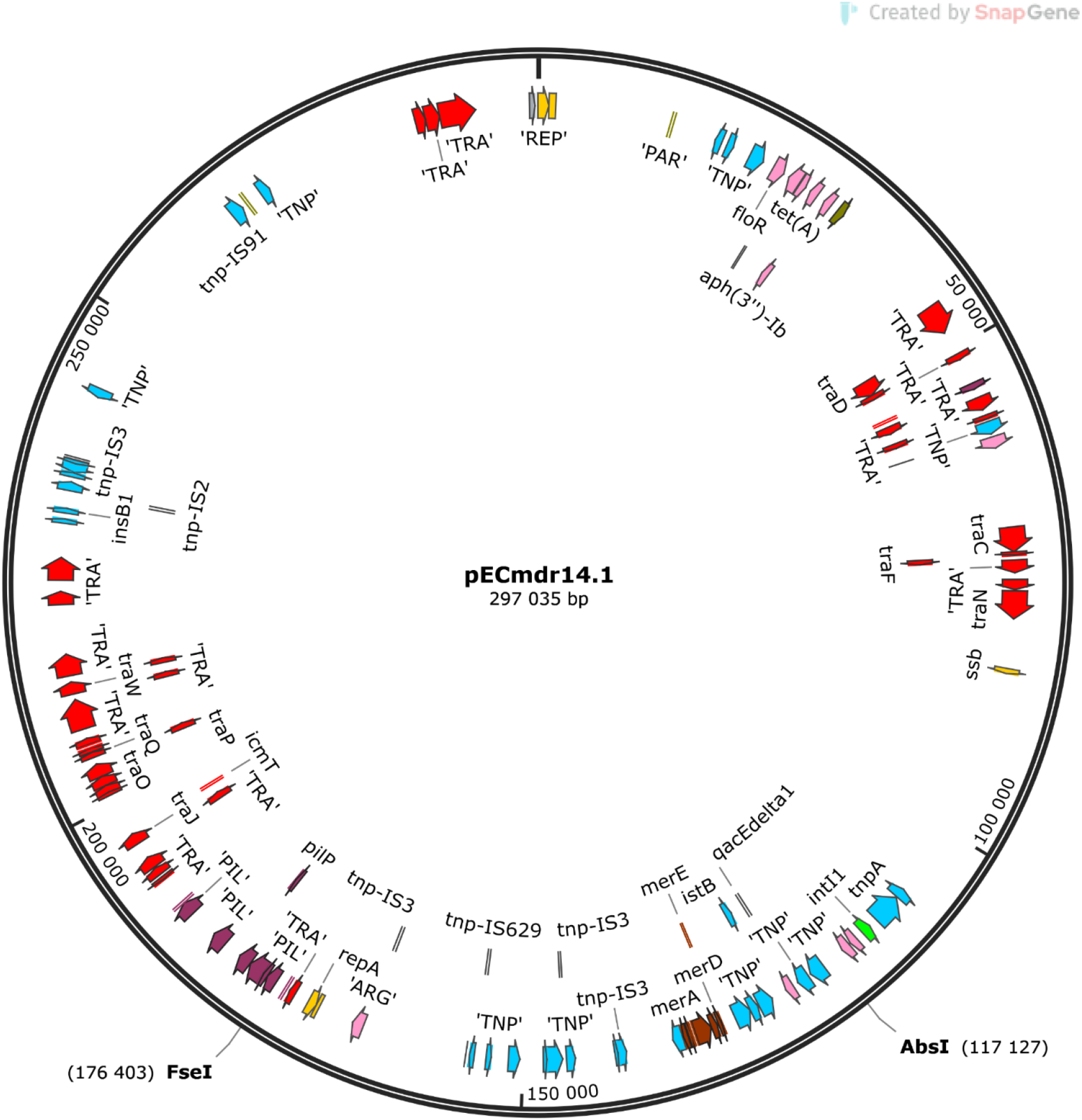
Linear and circular representation of pECmdr14.1 plasmid; pink-ARGs, red-conjugal transfer genes (TRA), blue-transposon transposase genes (TNP), brown-heavy metal resistance genes, yellow-replication system (REP), green-integron integrase, khaki-partitioning system (PAR), gray-mob system (MOB), plum-pilus system (PIL); a name given in quotation marks indicates that the gene function has been assigned based on the homology of the protein it encodes.

**Table 2.** ARGs, MGEs, integron integrase genes, transposon transposase genes, conjugal transfer system genes and heavy metal resistance genes.

| plasmid no. | size [bp] | ARGs | MGEs | integron<br>integrase gene | transposon<br>transposase<br>gene | conjugal transfer<br>system | heavy metal<br>resistance genes |
| --- | --- | --- | --- | --- | --- | --- | --- |
| pECmdr13.1 | 174994 | - | *cn_21731_IS2*<br>*cn_28888_IS2*<br>*cn_28883_ISEc27*<br>*cn_3769_ISEc27*<br>*cn_26991_IS629*<br>*cn_6274_ISSen9*<br>*cn_17058_ISSen9*<br>*ISEc12*<br>*ISEc8*<br>*IS421*<br>*IS5075*<br>*IS2*<br>*ISEc27*<br>*IS629*<br>*ISSen13*<br>*ISSen9* | - | *tnp-IS629*<br>*tnp-IS3*<br>*tnp-IS2*<br>*tnp-IS186B*<br>*tnp-IS91*<br>*insB1* | *traX*<br>*traI*<br>*traD*<br>*traS*<br>*traG*<br>*trbJ*<br>*trbB*<br>*traQ*<br>*trbA*<br>*trbF*<br>*traN*<br>*trbC*<br>*trbI*<br>*trbR*<br>*traV*<br>*trbG*<br>*traK*<br>*traE*<br>*traL*<br>*traY*<br>*traJ*<br>*traA* | - |
| *pECmdr13.2 | 158923 | *tet(A)*<br>*aph(6)-Id*<br>*aph(3'')-Ib*<br>*sul2*<br>*bla<sub>CMY-2</sub>*<br>*aadA1*<br>*aac(3)-Vla*<br>*sul1* | *cn_28214_IS5075*<br>*cn_2064_IS26*<br>*ISEc58*<br>*IS5075*<br>*IS26*<br>*Tn6196* | *intI1* | *tnpA*<br>*spoIVCA* | *traD*<br>*traL*<br>*traV*<br>*traN* | *merE*<br>*merD*<br>*merB*<br>*merA*<br>*merP*<br>*merT*<br>*merR* |
| *pECmdr13.3 | 45794 | *aph(3')-Ia*<br>*qnrS1*<br>*tet(A)*<br>*bla<sub>TEM-176</sub>* | *cn_22462_IS102*<br>*cn_3587_IS26*<br>*IS102*<br>*IS26*<br>*Tn2* | - | - | - | - |
| *pECmdr14.1 | 297035 | *floR*<br>*tet(A)*<br>*aph(6)-Id*<br>*aph(3'')-Ib*<br>*sul2*<br>*bla<sub>CMY-2</sub>*<br>*aadA1*<br>*aac(3)-Vla*<br>*sul1* | *cn_28214_IS5075*<br>*cn_3781_IS2*<br>*cn_16123_IS629*<br>*cn_6432_IS629*<br>*cn_2064_IS26*<br>*ISEc8*<br>*ISEc45*<br>*ISEc58*<br>*IS5075*<br>*IS2*<br>*IS629*<br>*ISVsa3*<br>*IS26*<br>*ISSen9*<br>*Tn6196* | *intI1* | *tnpA*<br>*spoIVCA*<br>*tnp-IS3*<br>*tnp-IS629*<br>*tnpB*<br>*insE*<br>*tnp-IS1*<br>*insB1*<br>*tnp-IS2*<br>*tnp-IS256*<br>*iS285*<br>*tnp-IS91* | *traD*<br>*traL*<br>*traV*<br>*traN*<br>*traO*<br>*traP*<br>*traQ*<br>*traW*<br>*traX* | *merE*<br>*merD*<br>*merB*<br>*merA*<br>*merP*<br>*merT*<br>*merR* |
| pECmdr14.2 | 157029 | - | *cn_21300_ISSen9*<br>*cn_10095_ISSen9*<br>*ISEc8*<br>*IS21*<br>*IS3*<br>*ISEc37*<br>*IS30*<br>*IS629*<br>*IS5075*<br>*ISSen13*<br>*ISSen9*<br>*ISSen9*<br>*ISKpn42* | - | *tnp-IS629*<br>*tnp-IS3*<br>*tnp-IS2*<br>*tnp-IS186B*<br>*tnp-IS91*<br>*insB1* | *traD*<br>*traG*<br>*trbJ*<br>*trbB*<br>*trbQ*<br>*trbA*<br>*traF*<br>*trbE*<br>*traN*<br>*trbC*<br>*traU*<br>*traW*<br>*trbI*<br>*traC*<br>*traR*<br>*traV*<br>*trbG*<br>*trbD*<br>*traK*<br>*traE*<br>*traL*<br>*traA*<br>*traY,*<br>*traJ*<br>*psiB* |  |
| pECmdr14.3 | 99355 | - | ISEc45 | - | - |  |  |

**cn** - composite transposon; **IS** - insertion sequence; **Tn** -transposon; \* - R-plasmids selected for analyses

Alignment of ARG-coding sequences and MGE allowed the identification of ARGs located within defined and described MGE, such as IS26, ISEcp1, or ‘ISclustersTn’, which were in turn situated on the plasmids identified during the study (Fig. 6A-G).

**Figure 6.**
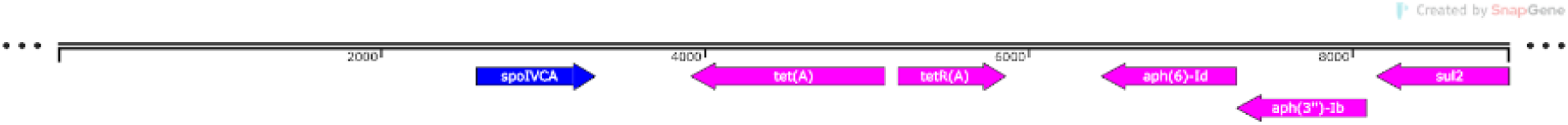
**A.** pECmdr13.2 IS26 (8965 bp) (pink-ARG, blue-transposon transposase genes TNP)

**Figure 6.**
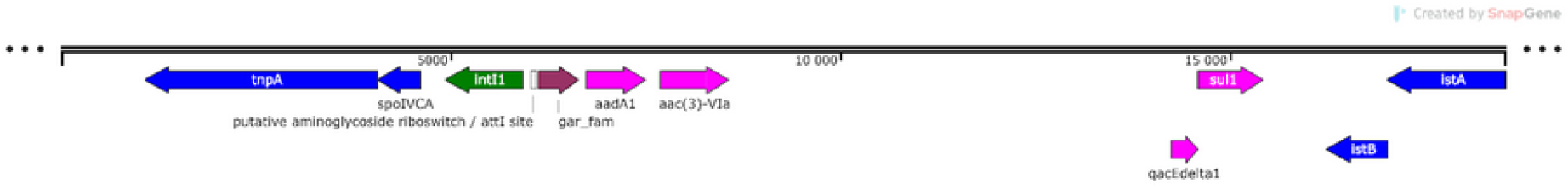
**B.** pECmdr13.2 ISclusterTn (18511 bp); pink-ARGs, blue-transposon transposase genes (TNP), green-integron integrase

**Figure 6.**
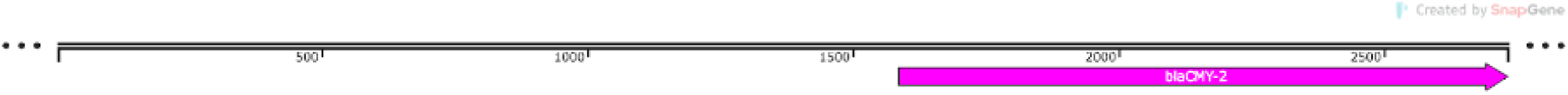
**C.** pECmdr13.2 ISEcp1 (2731 bp); pink-ARG

**Figure 6.**
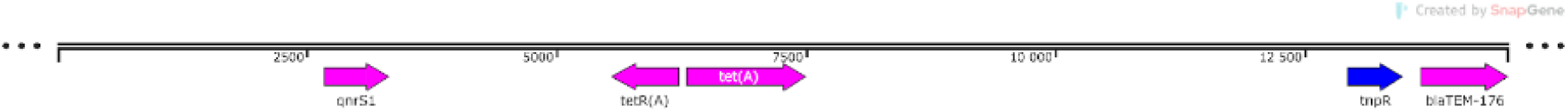
**D.** pECmdr13.3 ISclusterTn (14527 bp); pink-ARGs, blue-transposon transposase genes (TNP)

**Figure 6.**
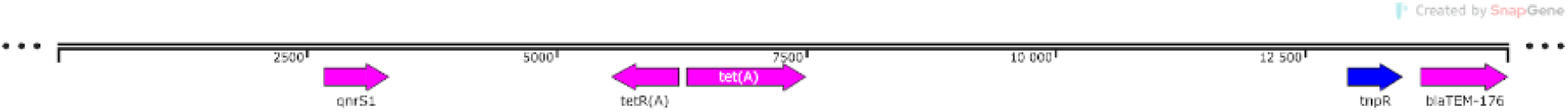
**E.** pECmdr14.1 ISclusterTn (14527); pink-ARGs, blue-transposon transposase genes (TNP)

**Figure 6.**
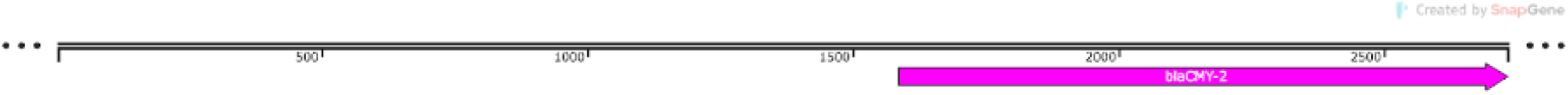
**F.** pECmdr14.1 ISEcp1 (2731 bp); pink-ARG

**Figure 6.**
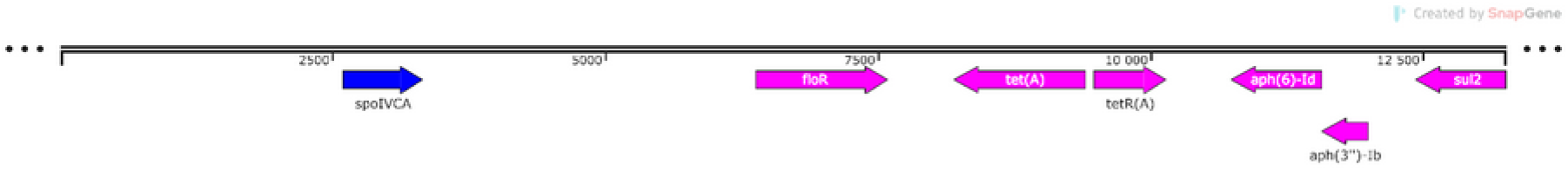
**G.** pECmdr14.1 IS26 (13249 bp); pink-ARGs, blue-transposon transposase genes (TNP)

A detailed search of the PLSDB provides data on the isolation source, biosample origin, country, and first report of similar plasmids (Table 3), as well as previously-reported plasmid host range (Table 4).

**Table 3.** Region, isolation source, and host biosample of similar plasmids (plasmids identity 0.99).

| Plasmid no. | Plasmid type | Firstly reported | Region | Isolation source | Host biosample |
| --- | --- | --- | --- | --- | --- |
| pECmdr13.2 | IncC_1<br>JN157804 | 2006 | France<br>USA<br>South Korea<br>Japan | river<br>urine<br>nose<br>stool<br>pus<br>abscess<br>peritoneal fluid<br>anal swab<br>abdominal drain fluid<br>nasal swab<br>tracheal secretion<br>bovine peripheral lymph node<br>bovine adipose trim<br>bovine hide<br>bovine pre-evisceration carcass<br>ground beef<br>retail meat<br>gut | Bovine<br>Human<br>Swine<br>Chicken (Gallus gallus)<br>Mouse<br>Snake<br>Turkey<br>Wild bird |
| pECmdr13.3 | IncX1_1<br>EU370913 | 2019 | France<br>Pakistan<br>India<br>USA<br>South Korea<br>Singapore<br>China<br>Laos | river<br>feces<br>urine<br>blood<br>chicken meat<br>wet market | Chicken<br>Human<br>Migratory bird |
| pECmdr14.1 | IncC_1<br>JN157804 | 2006 | France<br>USA<br>Mexico<br>Japan<br>South Korea<br>Germany<br>Taiwan<br>Thailand<br>India<br>China<br>Haiti<br>Switzerland<br>China<br>Myanmar | river<br>stool<br>retailed meat<br>bovine peripheral lymph node<br>ground beef<br>bovine adipose trim<br>abdominal<br>bovine pre-evisceration carcass<br>bovine hide<br>Peritoneal fluid<br>abdominal drain fluid<br>anal swab | Cattle<br>Human<br>Chicken (Gallus gallus)<br>Swine<br>Turkey<br>Wild bird |
|  |  |  | Canada | nasal swab<br>tracheal secretion<br>urine<br>mouse gut<br>diarrheal snake<br>pus<br>abscess<br>wound<br>slaughterhouse |  |

**Table 4.** The host range for similar plasmids (plasmid identity 0.99)

| Plasmid similar to: | Identified in the following bacteria: |
| --- | --- |
| pECmdr13.2 | *Salmonella enterica subsp. enterica*<br>*Escherichia coli*<br>*Proteus mirabilis*<br>*Vibrio alginolyticus*<br>*Klebsiella pneumoniae*<br>*Photobacterium damsela subsp. piscicida*<br>*Aeromonas salmonicida subsp. salmonicida*<br>*Citrobacter freundii*<br>*Aeromonas hydrophila* |
| pECmdr13.3 | *Escherichia coli*<br>*Klebsiella pneumoniae*<br>*Salmonella enterica subsp. enterica* |
| pECmdr14.1 | *Escherichia coli*<br>*Salmonella enterica subsp. enterica*<br>*Aeromonas salmonicida subsp. salmonicida*<br>*Klebsiella pneumoniae*<br>*Klebsiella quasipneumoniae*<br>*Proteus mirabilis*<br>*Aeromonas hydrophila*<br>*Vibrio cholerae*<br>*Shewanella algae*<br>*Vibrio alginolyticus*<br>*Citrobacter freundii*<br>*Edwardsiella ictaluri*<br>*Photobacterium damsela* |

## Discussion

Due to its presence in the veterinary, medical, and natural environment, *E. coli* is considered a specific vector for the transmission of ARGs, so-called acquired resistance (14, 15). The bacteria found in chicken manure or chicken litter are not limited to *E. coli*; they often co-isolate with other foodborne bacteria such as *Salmonella* and *Campylobacter* spp. (16–18). This is an important issue since such material is utilized as manure in agriculture globally (17, 19). These bacteria pose a serious risk of antimicrobial resistance transmission between animals, humans, and the environment, i.e. the three links of the One Health concept, and demonstrate considerable survival in water, soil, and crops: the lifespan of pathogenic bacteria ranges from several days to 10 years in soil and from several days to a year on plants (20, 21).

The main process responsible for disseminating ARGs is horizontal gene transfer (HGT) (22). The principal route of HGT in bacterial communities is most likely conjugation, and conjugation elements, such as conjugative plasmids, often harbor multiple ARGs (23). In addition, ARGs are often encoded on non-conjugative plasmids, transposons, integrative, and bacteriophages (24), whose presence facilitates HGT. HGT is most likely to occur when bacterial density is high, particularly where high numbers of bacteria are present in a given environment, and under stressing conditions, and selective pressure (14). HGT is also affected by genetic diversity in the environment: while high biodiversity may increase the pool of potentially exchangeable genes in the bacterial population, with HGT occuring even between distantly-related species, high diversity is considered a barrier to HGT (25).

Antimicrobial resistance is undoubtedly associated with resistance plasmids, which spread rapidly through conjugation. Accordingly, plasmids play a significant role in the worldwide dissemination of resistance, including MDR (26–28). Therefore, identifying and characterizing R-plasmids and their association with different bacterial hosts is key to understanding their involvement in the transfer of antimicrobial resistance. So far, little information exists on the fully-characterized plasmids identified in chicken manure isolates. A suitable example could be the aminoglycoside resistance plasmids (arr-3 and aacA) pRKZ3 (IncQ) and pKANJ7 (IncX), which were isolated from pig and chicken manure (29). The present study identified two IncC R-plasmids, pECmdr13.2 and pECmdr14.1, carrying genes for resistance to tetracyclines, aminoglycosides, β-lactams, sulfonamides, fluoroquinolones and to mercury; in addition, pECmdr14.1 demsontrated resistance to phenicol. The study also identified one IncX1 pECmdr13.3 R-plasmid, carrying genes for resistance to aminoglycosides, β-lactams, tetracyclines, and fluoroquinolones. So far, such plasmids have not been reported in Poland.

Incompatibility Group C (IncC) plasmids have a broad host range; the group can be divided into several types based on differences in sequence arising from homologous recombination events between type 1 and type 2 IncC backbones (30, 31). This variability also results from insertion and deletion events or inversion mechanisms involving IS26 (32). IncC plasmids occur widely in Gram-negative bacteria and are responsible for transmitting resistance to several different antibiotics, such as aminoglycoside and fluoroquinolone, as well as cephalosporins (*bla_CMY_* cephalosporinase genes) and carbapenems (*bla_NDM_* carbapenemase genes) which have considerable clinical significance (26, 31, 33, 34). In contrast, the IncX plasmids, and their subgroups (IncX1—IncX8), have a narrow spectrum of hosts, including *E. coli, Pseudomonas aeruginosa, Klebsiella pneumoniae* and *Salmonella enterica* (35).

These plasmids have been associated with antibiotic resistance to β-lactams, including carbapenems, aminoglycosides, quinolones, tetracyclines, streptomycin, and amphenicols, which are associated with mobile elements, such as insertion sequences, integrons and transposons (36). As shown in Table 2, the plasmids identified in this work have host ranges encompassing pathogenic and opportunistic bacteria capable of living in natural, veterinary, and clinical environments, which further emphasizes the importance of characterizing R-plasmids in these bacteria. Particularly noteworthy is the fact that most of these bacteria are categorized as critical and high-priority on the global priority pathogens list (37).

Previous studies found sulfonamide resistance gene (*sul1*), a tetracycline resistance gene (*tetA*), and three aminoglycoside resistance genes (*aadA*, *strA,* and *strB*) in plasmids isolated from chicken manure (35, 38). This is undoubtedly related to antibiotic therapy used in poultry farming: the most frequently-administered pharmaceuticals are beta-lactams, macrolides, polymyxins, quinolones, sulfonamides, and tetracyclines (39). There has been a surge of reports on multidrug-resistant *E. coli* strains isolated from chicken manure, resistant to β-lactams with ESBL (Extended-Spectrum Beta-Lactamase) and AmpC (clinically important β-lactamases; cephalosporinases) phenotypes, which are often also resistant to gentamicin (14). However, few studies have reported about plasmids coding for both antibiotic and metal resistance genes occurring in bacterial strains isolated from chicken feces. However, one study recent described the presence of ESBL/AmpC and mcr-5-carrying MDR plasmids isolated from *E. coli* and *K. pneumoniae* strains in Paraguayan poultry farms (40). IncHI2 plasmids identified in *E. coli* isolates from food-producing animals have also been found to carry the antibiotic resistance genes *bla_CTX-M_*/*oqxAB* with *aac (6′)-Ib-cr*, *floR*, *fosA3* and *rmtB*, as well as the heavy metal resistance genes *pco* and *sil* responsible for increasing the minimal inhibitory concentrations of CuSO4 and AgNO3 (41). Previously, similar IncHI2 plasmids carrying tetracycline, trimethoprim, and sulfonamide resistance genes and transposon Tn1696 related to mercury resistance were identified in two MDR *S. enterica* serovar Typhimurium isolates from Australian food-producing animals (42). Another study of *E. coli* strains from pig slaughterhouses in the United Kingdom showed various combinations of resistance to oxytetracycline, streptomycin, sulphonamide, ampicillin, chloramphenicol, trimethoprim–sulfamethoxazole, ceftiofur, amoxicillin–clavulanic acid, aztreonam, and nitrofurantoin, together with resistance to mercury, silver, or copper (*merA*, *merC,* and *pcoE* and s*ilA, silB*, and *silE* genes were detected) (43). A similar relationship was found in a 2024 study on the distribution and relationships of ARGs, heavy metal resistance genes, virulence factors and their transmission mechanisms of an MDR *E. coli* strain isolated from livestock manure and fertilized soil (44).

Therefore, the presence of metal resistance genes and ARGs on the identified plasmids supports their transmission by HGT and subsequent maintenance in the bacterial cell, even in the absence of selection pressure caused by antibiotics. This co-occurrence of genes specifying resistant phenotypes on one MGE is referred to as co-resistance (45, 46). Such genetic linkage between metal- and antibiotic-resistance traits has been reported on plasmids in *Enterobacteriaceae* and in other bacteria isolated from other animal feces, soil, or sewage (47–50). It has been shown that metals used in animal feed accumulate and persist in food animals, and may impact the development of AMR in primary animal and food production environments (51). Many studies indicate that the spread of AMR introduced through the application of manure to agricultural fields leads to its further spread within the food chain, and may pose a risk to human health (17, 52–54). Such a risk is underlined by the presence of plasmids carrying shared antibiotic and metal resistance in MDR strains of *E. coli*, a vector known to transmit AMR between One Health sectors: *E. coli* strains have been found to potentially transfer resistance between humans and chickens (55).

AMR continues to evolve and spread, with the main mechanism being HGT through plasmids. Therefore, the is a pressing need to identify and characterize R-plasmids and their relatnipships with different bacterial hosts to understand their involvement in the transfer of AMR determinants. The molecular identification of plasmid genotypes, the transposons and integrons located within them, or other insertion sequences involved in this process, would provide a clearer picture of the mechanism of AMR dissemination and its possible range. In turn, the characteristics of their hosts, i.e., the specific strains of bacteria, can improve the prediction of the risk to human and animal health. As such, our present findings fit perfectly into these goals.

## Conclusion

The study characterizes three *E. coli* plasmids, each carrying ARGs and various insertion elements (IS), two transposons (*Tn2* in pECmdr13.2 and *Tn6196* in pECmdr13.2 and pECmdr14.1) and a class 1 integron-integrase gene (intI1 n pECmdr13.2 and pECmdr14.1). The plasmids were found in two representative MDR isolates from chicken manure. No such plasmids have been previously reported in Poland, though similar ones have been identified in crucial human pathogens globally, and in natural settings like soil and water. Simialr plasmids to pECmdr13.2, pECmdr13.3, and pECmdr14.1, are present in various environments, clinical isolates, and farm animals including cattle, swine, and poultry.

The studied *E. coli* plasmids, identified in chicken manure, confer resistance to β-lactam antibiotics, including penicillins, even with β-lactamase inhibitors like clavulanic acid (*bla_CMY-2_*, *bla_TEM-176_),* second-to fourth-generation cephalosporins, aminoglycosides (coded for by several ARGs: *aph, acc* and *aad* families), tetracyclines (*tet[A]*), sulfonamide (*sul1*, *sul2*), fluoroquinolones (*qnrS1*) and phenicol (*floR*).

Notably, the *E. coli* plasmids were found to contain IS26 insertion elements and the intI1-integrase gene, associated with ARGs, which are common in other resistance plasmids isolated from key pathogens. Plasmids pECmdr13.2 and pECmdr14.1 also possess a mercury resistance operon, related to transposon Tn1696, promoting their maintenance through co-selection even without antibiotic pressure. All identified resistance plasmids carry conjugal transfer genes, enabling HGT.

The identified plasmids belong to incompatibility groups IncX1 and IncC; the IncX1 plasmids have a narrow host range (*Enterobacteriaceae* and *Pseudomonas*), while the IncC plasmids have a broad host range.

## Materials and Methods

### Isolation of *E. coli* strains

Chicken waste was collected from a commercial chicken meat production facility located in the central part of Poland (Masovian district). The samples were taken from two points in the production chain: CM—chicken manure from laying hens and CL—chicken litter from broilers. Sampling details were described previously by Błażejewska et. al (17). Samples of chicken litter and manure (0.1 kg per sampling point) were collected from 10 locations for each waste type: the former in the chicken coop and the latter under the cages. The samples were then divided by type and pooled into one representative sample for each type for further analysis. All polled collections were taken in triplicate. The farm owners agreed to manure sampling.

The collected samples were transported to the laboratory in a refrigerator, stored at 4 °C, and processed within 24 hours. To isolate antibiotic-resistant *E. coli* strains, the manure sample was first enriched in Luria Bertani broth by adding 1 g of feces to 9 mL of liquid medium. The cultures were incubated at 30 °C and 37 °C for 24 hours and 48 hours. After incubation, the bacterial suspensions were diluted (10^−1^, 10^−2^, and 10^−3^) and 100 µL of undiluted sample and each dilution were plated on Eosin Methylene Blue Agar (Biomaxima, Lublin, Poland) and MacConkey Agar (Biomaxima, Lublin, Poland) supplemented with imipenem (16 mg/L) (Merck, Darmstadt, Germany) to search for carbapenem resistance phenotype; the same dilutions were also asdded to Eosin Methylene Blue Agar and MacConkey Agar both supplemented with cefotaxime (4 mg/L) (Merck, Darmstadt, Germany) for the Extended-spectrum beta-lactamase phenotype (37). Each isolation was performed as three biological replicates, followed by three technical ones.

The plates were incubated for 24 hours at 37 °C to select animal pathogens and 30 °C for environmental strains. Next, 24 to 48 colonies showing the morphology of the intended bacteria were taken and subjected to three consecutive streaks to get pure colonies. The pure cultures were stored in PBS/glycerol stocks (20% v/v) for further analysis.

### The susceptibility profiles of *E. coli* strains

The antibiotic susceptibility profiles of isolated strains were determined by The Kirby–Bauer test (56). For phenotype characterization, the following antimicrobial discs were used: IMP (10 µg), CIP (5 µg), CTX (5 µg), and CN (10 µg) (OXOID, USA). The inhibition zones were measured according to EUCAST recommendations, i.e. after incubation for 18 hours at 37 °C. Strains with different antimicrobial susceptibility profiles, indicated by the bacterial growth inhibition zone differing by approximately +/− 2 mm in diameter around at least one antibiotic disk, were considered non-repetitive and chosen for further research. The presence of *E. coli* was confirmed by MALDI-TOF MS/MS (matrix-assisted laser desorption/ionization system equipped with a time-of-flight mass spectrometer) (57).

The analyses were performed in external medical laboratory (ALAB Laboratoria Sp. z o. o., Poland), according to a standard diagnostic procedure. Identification was performed by aligning the peaks to the best matching reference data. The resulting log score was classified as follows: ≥2.3, highly probable species; between 2.0 and 2.3, certain genus and probable species; between 1.7 and 2.0, probable genus; and <1.7, non-reliable identification. Next, antibiotic susceptibility testing was performed with Vitek2 Compact equipment (BioMerieux, France) (58). Based on the results, strains were classified as sensitive (S), resistant (R), or intermediate (I). The AST-N331 susceptibility card was used.

Strains with the same resistance profile and origin were considered clones. The Plasmid profile of non-repetitive strains was determined in three ways: 1) Plasmid DNA was isolated with a commercially-available Plasmid Mini kit (A&A Biotechnology, Poland) 2) rapid alkaline lysis for the isolation of plasmid DNA (59), 3) plasmid visualization by Eckhardt electrophoresis (60). Plasmid DNA with similar profiles was additionally differentiated by digestion with the BamHI and HindIII restriction enzymes (NEB, USA). The digestion was performed according to the manufacturer’s recommendation. Strains with different profiles were chosen for sequencing.

### DNA extraction and sequencing

Genomic DNA from bacterial strains was isolated using a Genomic Mini kit (A&A Biotechnology, Gdynia, Poland). The DNA concentration was measured using a Qubit fluorometer and a dsDNA High Sensitivity Assay Kit (Thermo Fisher Scientific, USA), and purity was determined by measuring the A260/A280 absorbance ratio with a NanoDrop spectrophotometer (Thermo Fisher, USA). Only samples with concentrations higher than 10 ng/µL and an A260/A280 ratio ranging from 1.8 to 2.0 were analyzed. DNA samples were stored at −20 °C for further use. The DNA samples were isolated in triplicate.

### Genomic DNA sequencing on Oxford Nanopore MinION

Libraries were prepared from 500 ng of DNA samples after mechanical fragmentation by the syringe-based method (0.4 x 20 mm needle, 1 ml glass syringe, 200 µl) and purification by Solid Phase Reversible Immobilization with Kapa Pure Beads (Roche, cat. no. 07983298001, elution for 10 minutes at 37°C). Native Barcoding method was applied (Oxford Nanopore, cat. no. SQK-NBD112.24, Version: NBE_9134_v112_revE_01Dec2021). The NanoPore MINIon MkB1 R10.4 flow cell and reagents (cat. no. FLO-MIN112) was used for sequencing with standard procedures. Guppy software (version 6.1.2, -- flowcell FLO-MIN112 --kit SQK-NBD112-24 options) was applied for basecalling.

### Genomic DNA sequencing on Illumina NovaSeq 6000

Briefly, 100 ng of genomic DNA was fragmented to 300 bp by ultrasonication (S220 Covaris, Duty Factor: 10%, Peak Incident Power: 140, Cycles per Burst: 200, Treatment Time: 80s) and subjected to library construction with the KAPA Hyper Prep Kit (Roche, cat. no. 07962363001), according to the manufacturer’s protocol, with five cycles of amplification; however, TruSeq DNA UD Indexes adapters were used (Illumina, cat. no. 20020590). Libraries were size-selected in a two-step SPRI method and with a sample-to-reagent volume ratio of 1) step - 1: 0.75, 2) step - 1:0.85 using Kapa HyperPure Beads (Roche, cat. no. 07983298001). Sequencing was performed using Pair-end 2×100 cycle mode on the Illumina NovaSeq 6000 system (NovaSeq 6000 S1 Reagent Kit v1.5 200 cycles Illumina, cat. no. 20028317; 0.5% of the PhiX control library, Illumina, cat. no. FC-110-3001) and the standard clustering procedure.

### Genome assembly and data analysis

Initial sequence quality metrics for Illumina data were obtained using FASTQC v.0.12.0 (http://www.bioinformatics.babraham.ac.uk/projects/fastqc/) (61) and subjected to quality trimming using fastp v.0.23.2 (62). Nanopore reads were quality filtered with NanoFilt v.2.8.0, discarding reads below 1kb and QV<12 (63), and residual adapters were removed using Porechop v.0.2.4 (https://github.com/rrwick/Porechop). The quality of the final long-read dataset was evaluated using NanoPlot v. 1.41.6 (63).

Long-read assembly was performed using Trycycler v.0.5.3 pipeline (64). Nanopore reads were initially assembled using four long-read assemblers: Flye v2.9, Unicycler v0.4.8, Raven v1.8.1 and Miniasm v0.3-r179. The resulting assemblies were then reconciled and circularized, and a consensus sequence was generated, which underwent further polishing with medaka_consensus (https://github.com/nanoporetech/medaka). The long-read assembled contigs were further polished with short Illumina reads using polypolish (https://github.com/rrwick/Polypolish) v. 0.5.0 (64) and POLCA v. 4.0.5 (65). Finally, all of the sequence errors and misassemblies were manually corrected using SeqMan software v. 9.1 (DNAStar) to obtain a complete nucleotide sequence for the bacterial genome.

The annotation of consensus sequences was performed using Bakta (66). Plasmid replicon sequences were searched against the PLSDB database (https://ccb-microbe.cs.uni-saarland.de/plsdb/). Antimicrobial resistance genes were identified and localized in the genomic sequences using abricate (https://github.com/tseemann/abricate). Mobile genetic element detection was performed using mobileelementfinder (67), and the ARG-associated mobilome was characterized using VRprofile2 (68).

The plasmid sequences were visualized using SnapGene Viewer software (SnapGene® software from Dotmatics; available at snapgene.com).

## Author contributions

M.Z. – Methodology, Formal analysis, Investigation, Data Curation, Writing – Original Draft, Visualization; A.B. – Formal analysis, Investigation; J.G. – Formal analysis, Data Curation; A.D., S.M., K.G – Sequencing and primary Data Curation; P.K – Investigation, Data Curation; M.P. – Conceptualization, Writing – Review and Editing, Project administration, Funding acquisition.

## Acknowledgments

The research was funded by National Science Centre, Poland (2017/25/Z/NZ7/03026), grant under the European Horizon 2020, in the frame of the JPI-EC-AMR Joint Transnational Call (JPIAMR), JPI-EC-AMR JTC 2017, project INART–“Intervention of antibiotic resistance transfer into the food chain” to MP and partially in the frame of the “Excellence Initiative—Research University (2020–2026)” Program at the University of Warsaw. Sequencing was performed in GCF CeNT UW (RRID:SCR_022718), using NovaSeq 6000 platform financed by Polish Ministry of Science and Higher Education (decision no. 6817/IA/SP/2018 of 2018-04-10).

## Data availability

The genome sequences of *E. coli* strains generated during this study have been deposited in GenBank (NCBI) with the accession number PRJNA942482. Plasmids datasets generated during this study have been deposited in the University of Warsaw repository with the doi number https://doi.org/10.58132/8ADBIC.

